# Anatomical and functional organization of cardiac fibers in the porcine cervical vagus nerve allows spatially selective efferent neuromodulation

**DOI:** 10.1101/2024.01.09.574861

**Authors:** Nicole Thompson, Enrico Ravagli, Svetlana Mastitskaya, Ronald Challita, Joseph Hadaya, Francesco Iacoviello, Ahmad Shah Idil, Paul R. Shearing, Olujimi A. Ajijola, Jeffrey L. Ardell, Kalyanam Shivkumar, David Holder, Kirill Aristovich

**Affiliations:** EIT and Neurophysiology Research Group, Department of Medical Physics and Biomedical Engineering, University College London, London, United Kingdom; UCLA Cardiac Arrhythmia Center and Neurocardiology Research Program of Excellence, David Geffen School of Medicine at UCLA, Los Angeles, California, USA; Electrochemical Innovation Lab, Department of Chemical Engineering, University College London, London, United Kingdom

**Author notes:** Address for correspondence: Dr Nicole Thompson, EIT and Neurophysiology Research Group, Department of Medical Physics and Biomedical Engineering, University College London, London, United Kingdom. Twitter handle: @nog_thompson. ORCID: https://orcid.org/0000-0002-4153-9614. Joint first authors (Nicole Thompson, Enrico Ravagli).

**Keywords:** vagus nerve stimulation (VNS), spatially-selective, neuromodulation, cardiac, afferent and efferent, heart failure

## Abstract

Cardiac disease progression reflects the dynamic interaction between adversely remodeled neurohumoral control systems and an abnormal cardiac substrate. Vagal nerve stimulation (VNS) is an attractive neuromodulatory option to dampen this dynamic interaction; however, it is limited by off-target effects. Spatially-selective VNS (sVNS) offers a promising solution to induce cardioprotection while mitigating off-target effects by specifically targeting pre-ganglionic parasympathetic efferent cardiac fibers. This approach also has the potential to enhance therapeutic outcomes by eliminating time-consuming titration required for optimal VNS. Recent studies have demonstrated the independent modulation of breathing rate, heart rate, and laryngeal contraction through sVNS. However, the spatial organization of afferent and efferent cardiac-related fibers within the vagus nerve remains unexplored.

By using trial-and-error sVNS *in vivo* in combination with *ex vivo* micro-computed tomography fascicle tracing, we show the significant spatial separation of cardiac afferent and efferent fibers (179±55° SD microCT, p<0.05 and 200±137° SD, p<0.05 sVNS – degrees of separation across a cross-section of nerve) at the mid-cervical level. We also show that cardiac afferent fibers are located in proximity to pulmonary fibers consistent with recent findings of cardiopulmonary convergent neurons and circuits. We demonstrate the ability of sVNS to selectively elicit desired scalable heart rate decrease without stimulating afferent-related reflexes.

By elucidating the spatial organization of cardiac-related fibers within the vagus nerve, our findings pave the way for more targeted neuromodulation, thereby reducing off-target effects and eliminating the need for titration. This, in turn, will enhance the precision and efficacy of VNS therapy in treating cardiac pathology, allowing for improved therapeutic efficacy.

**Condensed Abstract:** Spatially-selective vagus nerve stimulation (sVNS) presents a promising approach for addressing chronic heart disease with enhanced precision. Our study reveals significant spatial separation between cardiac afferent and efferent fibers in the vagus nerve, particularly at the mid-cervical level. Utilizing trial-and-error sVNS in vivo and micro-computed tomography fascicle tracing, we demonstrate the potential for targeted neuromodulation, achieving therapeutic effects like scalable heart rate decrease without stimulating afferent-related reflexes. This spatial understanding opens avenues for more effective VNS therapy, minimizing off-target effects and eliminating the need for titration, thereby expediting therapeutic outcomes in myocardial infarction and related conditions.

**Tweet:** With functional and structural imaging, we found organization of vagal efferent & afferent cardiac regions. We can selectively activate only cardiac efferents to achieve bradycardia; desired to reduce the effects of sympathetic overactivation associated with heart disease #VNS #Cardiac #VagusNerve

**Key Points:**

- Spatially-selective vagus nerve stimulation (sVNS) presents a promising approach for addressing chronic heart disease with enhanced precision.
- Our study reveals significant spatial separation between cardiac afferent and efferent fibers in the vagus nerve, particularly at the mid-cervical level.
- Utilizing trial-and-error sVNS in vivo and micro-computed tomography fascicle tracing, we demonstrate the potential for targeted neuromodulation, achieving therapeutic effects like scalable heart rate decrease without stimulating afferent-related reflexes.
- This spatial understanding opens avenues for more effective VNS therapy, minimizing off-target effects and eliminating the need for titration, thereby expediting therapeutic outcomes in myocardial infarction and related conditions.

## Introduction

The vagus nerve is a major focus of neuromodulation due to its extensive distribution and complex neural pathways to and from the heart, larynx, lungs, and abdominal viscera. It plays a vital role in regulating various bodily functions. As the main peripheral nerve of the parasympathetic branch of the autonomic nervous system, it controls heart rate, blood pressure, digestion, respiration, and immunity. By applying electrical stimulation to the nerve, these physiological effects can either be induced or inhibited. At present, vagus nerve stimulation (VNS) is used to treat drug-resistant depression and epilepsy, but a particularly critical application for VNS is in cardiac pathologies such as heart failure (HF), ischemic heart disease, myocardial infarction (MI), hypertension and arrhythmia ^1–3^.

The autonomic nervous system (ANS), via its vagal and sympathetic limbs, controls cardiac contractile and electrophysiologic function. Following MI, ANS dysfunction results in excessive reflex activation of efferent sympathetic tone to the heart to compensate for the reduced functional capacity of the damaged heart. MI further leads to downregulation of parasympathetic (vagal) activity, this is associated with lethal ventricular arrhythmias ^4^ and is intricately involved in the process of maladaptive cardiac remodeling ^5^ that ultimately leads to cardiac mortality. By performing VNS, regional cardiac electrical and mechanical function is maintained, susceptibility of the myocardium to harmful arrhythmias and the effects of sympathetic system overactivation can be reduced thereby mitigating heart disease progression ^3,6^. In preclinical studies, reactive VNS has preserved autonomic control and induced cardioprotection when applied early on post-MI ^3^. VNS has also shown promise for treating HF in recent clinical trials where improvements from baseline were observed in standard deviation in normal-to-normal R-R intervals, left ventricular ejection fraction, Minnesota Living with HF mean score, and an improvement in 6-min walk distance was observed ^7,8^.

However, there are two major problems with clinical applications of VNS. First, the current paradigm of VNS is to stimulate the entire nerve with the FDA approved implantable VNS interventions that lack spatial selectivity capabilities, leading to indiscriminate stimulation of non-targeted effectors and undesired side effects ^9^. These off-target effects are largely due to afferent fiber activation; although some side effects such as pain which could be argued to be attributed to efferent activation, with pain reported during VNS in less than 3% of patients in the ANTHEM HF with reduced ejection fraction (HFrEF) trial ^7,8^. Secondly, due primarily to these off-target effects on respiratory and GI function, VNS on the whole nerve takes time (∼4 weeks) to titrate to therapeutic levels ^10,11^. Given that time is critical following MI, with only adverse outcomes expected from delays due to the irreversible nature of many post-MI neural and cardiac events ^12–18^, it is advantageous to apply VNS during or soon after a heart attack, thereby reducing infarct size, preventing ANS dysfunction and improving survival.

A possible solution to overcome these hurdles is the use of spatially-selective VNS (sVNS). Spatially-selective VNS was first demonstrated in its modern form in the vagus nerve of sheep ^19^ for evoking separate respiratory and cardiac effects. Briefly, this work used an electrode array geometry consisting of 14 electrode pairs, placed at equally spaced angular positions around the circumference of the nerve over 2 electrode rings. More recently, sVNS of the vagus nerve was also reported in the pig vagus nerve using the same 14-electrode double-ring array ^20^, an 8-electrode single-ring array ^21^ and a 6-electrode FDA investigational device ^22^. Selective stimulation of the right vagus nerve with multiple contacts was also achieved recently for modulation of heart rate ^23^; however, this work employed an invasive Transversal Intrafascicular Multichannel Electrode (TIME) array. Recently, the organization of the cervical vagus nerve in swine was investigated by Thompson *et al* (2023) ^20^, who showed a degree of cross-sectional organization for branches innervating the larynx, heart and lungs, which cross-correlated between sVNS, micro-computed tomography (microCT) and electrical impedance tomography (EIT). Similar conclusions were drawn in another recent swine study ^21^. However, a deeper understanding of the functional and anatomical organization of cardiac nerve fibers in the cervical vagus is needed to perform sVNS to treat ischemic and non-ischemic heart disease.

Neural control of cardiac function involves a multi-tiered hierarchy of interdependent reflexes, and whole nerve VNS, when delivered at the cervical level, activates both the ascending and descending projections, engaging multiple levels of the cardiac neuroaxis ^2^. A key to effectively and rapidly treating heart disease and achieve therapeutic efficacy is stimulating the efferent vagal fibers whilst simultaneously avoiding activating vagal afferents. With whole nerve VNS stimulation, the titration to mitigate off target effects (e.g. cough and GI discomfort), takes 3-4 weeks ^10^. While this reactive temporal constraint is reasonable for stage 3 HFrEF patients ^7,8,24^, the adverse dynamic remodeling that occurs in neural and cardiac tissues post-MI is rapid and, in many cases, irreversible ^4,15–17^. As such, refinements in reactive VNS neuromodulation that shorten time to therapeutic efficacy increase the potential for cardioprotection ^3,25^.

Vagal efferent fibers regulate chronotropic, dromotropic, inotropic and lusitropic function of the heart by direct projections to heart tissues and by interacting with sympathetic projections to the heart ^26^. However, the location and organization of these respective fiber groups within the vagus nerve is unknown.

In this study, the organization of cardiac fibers in the mid-cervical porcine right vagus nerve was investigated by means of *in vivo* neuromodulation protocols and post-mortem imaging methods. The specific aims of the study were to answer the following questions:

1. Is there spatial separation and organization of efferent and afferent cardiac fibers over the cross section of the vagus nerve at the mid-cervical level?
2. Is it possible to perform sVNS of efferent cardiac fibers whilst avoiding activation of afferent fibers, which would allow for improved treatment of heart disease by reducing side effects and the need for titration?

## Methods

### 2.1 Experimental Design

The study was performed using Yorkshire castrated male pigs, weighing 53.7±3.7 kg. All experimental procedures were ethically reviewed by the UK Home Office and the Animal Welfare and Ethical Review Body (AWERB) and carried out in accordance with Animals (Scientific Procedures) Act 1986. Animal experiments, performed at UCLA, were approved by the UCLA Institutional Animal Care and Use Committee and performed in accordance with the National Institutes of Health’s Guide for the Care and Use of Laboratory Animals.

The study was designed with the aim of investigating the function and anatomy of the right cervical vagus and was comprised an *in vivo* and *ex vivo* phase. During the *in vivo* phase, sVNS of the right vagus nerve was performed with the nerve intact to assess the effect of both afferent and efferent pathways on cardiac function; sVNS was then repeated after RV distal vagotomy and left vagus vagotomy to isolate in steps the contribution of different neural pathways.

Additional electrophysiological recordings, namely sVNS of laryngeal and pulmonary function, were performed in the animals to collect more information about the functional anatomy of the cardiac branch in relation to other branches and fascicles.

Following completion of the electrophysiological recordings, animals were sacrificed, and RV were excised and subject to microCT imaging and organ-to-cervical branch tracing.

The full experimental protocol can be summarized as follows. Initially, the animal underwent anesthesia, followed by surgical exposure of both the right and left vagus nerves, with the placement of the sVNS cuff on the right vagus nerve (Figure 1). Subsequently, a bipolar stimulation electrode was carefully positioned on the recurrent laryngeal (RL) branch. During spontaneous breathing without ventilation, sVNS was conducted while monitoring EtCO2 and breathing rate to pinpoint the pulmonary branch. Electromyographic (EMG) electrodes were strategically inserted into the right recurrent laryngeal (RL) muscle region to facilitate the identification of the RL branch through sVNS. Next, sVNS was employed to assess cardiac function. Following these initial procedures, a right vagotomy was performed distal to the sVNS cuff, followed by a repeat of sVNS to evaluate cardiac function post-vagotomy. Finally, a left vagotomy was performed, and once again, sVNS of cardiac function was repeated post-vagotomy to conclude the experimental protocol.

**Figure 1.**
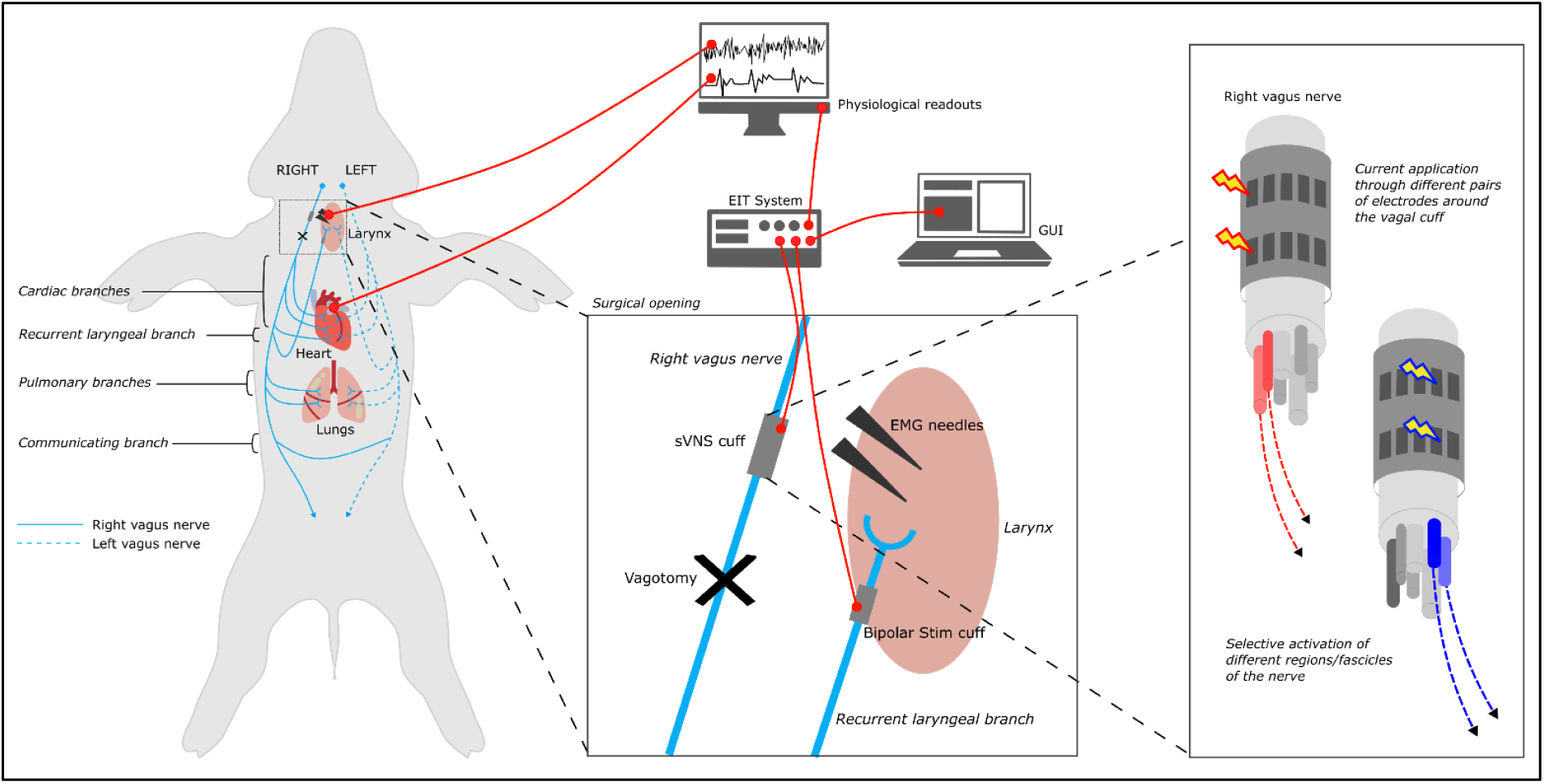
Experimental *in vivo* setup. A multi-electrode cuff for sVNS was applied to the right vagus. A custom electronic system performs sVNS on the cuff. A bipolar stimulation cuff is placed on the recurrent laryngeal branch and used to evoke neural traffic for recurrent laryngeal EIT. EMG needles are placed in the larynx muscle to detect contraction during recurrent laryngeal sVNS. A physiological readout system monitors blood pressure, ECG, respiration and other vital signs.

### 2.2 Surgery and anesthesia

On the day of the experiment, Tiletamine-Zolazepam (4-6mg/kg, intramuscular) was used for induction of anesthesia. Animals were endotracheally intubated and mechanically ventilated (tidal volume 400 to 600 mL, rate 12-16 breaths/minute). Anesthesia was then maintained with isoflurane vaporized (1-4% inhaled). A continuous rate infusion of fentanyl (0.2 μg/kg/min) was started after induction and continued during subsequent surgical procedures for analgesia.

After induction of general anesthesia, the animal was positioned in dorsal recumbency. Indwelling catheters were percutaneously placed using ultrasound guidance in both femoral arteries (for blood pressure and blood gas monitoring) and femoral veins. The animal was instrumented with a 12-lead electrocardiogram (ECG) and a pulse oximeter. A spirometer was connected to the tracheal tube. The animal was mechanically ventilated using pressure control mode for the duration of the surgery and for most of the experiment, except when sVNS were applied for identification of pulmonary responses. Between these periods, if required, animals were placed onto mechanical ventilation to restore normal levels of CO_2_ (between 35−45 mmHg). Body temperature was maintained using a hot air warming system if necessary. Lactate ringer fluid therapy at a rate of 5 ml/kg/h was administered intravenously throughout the procedure.

Following completion of surgical procedures, anesthesia was transitioned from isoflurane to alpha-chloralose (25-50 mg/kg/hour, intravenous), an anesthetic that does not depress cardiac and autonomic reflexes.

Routine anesthesia monitoring included vital parameters such as ECG and invasive arterial blood pressure, central venous pressure, end-tidal CO2 (EtCO2), end-tidal isoflurane (Et Iso), pulse oximetry and core body temperature (via rectal probe). The anesthetized animals were monitored to maintain physiological parameters within the normal limits for the species, specifically: heart rate (HR): 80-120 bpm; mean blood pressure: 70 -120mmHg; EtCO2: 35-45mmHg; and bispectral index: 50-70. Some parameters (arterial blood pressure, central venous pressure, ECG, EtCO2, Et Iso) were also digitally recorded using a 16 channel PowerLab acquisition system (ADInstruments) with LabChart 8 software at 2 kHz sampling frequency. Levels of anesthetic were adjusted accordingly by the anesthetist.

The ventral neck region was clipped and aseptically prepared using chlorhexidine-based solutions. Longitudinal 20 cm skin incisions were made using monopolar electrocautery centered immediately to the left and right of the trachea, respectively. The incision was continued through the subcutaneous tissue and the sternohyoideus musculature using a sharp/blunt technique until encountering the carotid sheath and vagus nerve. A 5-7 cm long segment of the left and right vagus nerves were circumferentially isolated by blunt dissection to allow placement of a sVNS electrode. The electrode cuff was placed around each nerve by carefully opening the cuff and sliding the vagus inside it, with the cuff opening facing ventrally. The sVNS cuffs were placed at the mid-cervical level, approximately 2-3 cm from the nodose ganglion, respectively, keeping as consistent placement of the cuffs between animals as possible. To secure the cuff in place around the nerve during the experiment, the sutures incorporated into the design for opening the cuff were tied around the cuff circumference ensuring a tight fit. Electrical ground and earth electrodes were inserted into the surgical field. The impedances of the electrodes were <1 kOhm at 1 kHz.

The left and right recurrent laryngeal nerves were identified within the surgical field, and bipolar stimulating electrodes (CorTec GmbH, I.D. 1.2-2 mm) were placed around them. EMG needles were implanted into laryngeal muscle on each side, specifically the cricoarytenoid and thyroarytenoid muscles, to record laryngeal effects of sVNS. The surgical field was then rinsed with sterile saline.

After the initial round of sVNS for identification of pulmonary fascicles, the animal was put back on mechanical ventilation and anesthesia was switched to α-chloralose (50 mg/kg initial bolus, thereafter 20–50 mg·kg−1·h−1 i.v.). During a stabilization period (30 min) after anesthesia transition, rounds of sVNS were performed to identify the recurrent laryngeal fascicles by observing and recording laryngeal EMG response during stimulation. Following this step, and still during the stabilization period, EIT of evoked activity from recurrent laryngeal stimulation was performed. Afterwards, selective stimulation for identification of cardiac branches was performed, and the remaining 2-3 hours of the experiment were dedicated to spontaneous EIT recordings. A right vagotomy was performed approximately 1-2cm distal to the sVNS cuff and after another stabilization period, further sVNS performed on the right vagus nerve. Subsequently, a left vagotomy was performed 1-2cm proximal to the cuff on the left vagus nerve, followed by further sVNS. At the end of all recordings, an overdose of sodium pentobarbital (100 mg/kg i.v.) followed by saturated KCl (70-150 mg/kg i.v.) was used for euthanasia.

### 2.3 Instrumentation for sVNS

All sVNS recordings in this study were performed using the ScouseTom EIT system ^27^.

This system integrates a commercial grade electroencephalogram (EEG) amplifier (Actichamp, Brain Products GmbH, Germany), two benchtop current sources (Keithley 6221 AC, Tektronix, United Kingdom) and custom PCB boards for multiplexing, timing and communication. The EEG amplifier has 24-bit resolution (±400 mV range, internal gain =5), an internal 10kHz hardware anti-aliasing filter, and is able to sample data in a true parallel configuration at 50 kSamples/s. Auxiliary channels from the amplifier (±5 V range, internal gain =1), operating at the same sampling frequency and resolution, were used to acquire physiological signals ECG, EtCO2, laryngeal EMG, systemic and ventricular blood pressure from the clinical monitoring system.

While this system was developed mainly for neural EIT, it can also be used for performing sVNS due to the compatible hardware features. In EIT mode, the custom multiplexer board in the system is set up to route low-amplitude sinusoidal current from one of the current sources to the nerve cuff electrode. The second current source is used to deliver current pulses to a nerve branch if the system is used for evoked EIT, or left unused if EIT of spontaneous neural traffic is being performed. In sVNS mode, the pulse-generating source is connected directly to the multiplexer board to perform sVNS, while the sinusoidal source is left on standby. For stimulation amplitudes higher than 2mA, the current source was routed directly to the electrode array to avoid cross-talk between channels on the multiplexer board.

### 2.4 Spatially-selective Vagus Nerve Stimulation

Spatially-selective stimulation of the right vagus nerve was performed using electrode arrays embedded within circular epineural nerve cuffs.

Electrode arrays had the same geometry as the ones used by Aristovich *et al* (2021) ^19^ and Thompson *et al* (2023) ^20^, and were manufactured using the same process ^28^. Briefly, arrays were composed of stainless-steel tracks contained in-between isolating layers of medical-grade silicone. Exposed pads were laser-patterned and coated with PEDOT:pTS to reduce contact impedance and noise from the electrode–electrolyte interface. Cuffs were glued to inner sides of silicone tubing to maintain tubular shape (2.7 mm inner diameter) for wrapping around the nerve. Arrays comprised two rings of 14 pads, each pad sized 3.00 × 0.35 mm^2^. Electrodes were spaced 0.32 mm along the circumference, edge-to-edge, and rings were spaced 3.1 mm between internal edges. Two external electrodes comprising the entire cross section of the nerve were present at the extremities and used as a control to perform stimulation of the full vagus nerve. These external electrodes were 0.46 mm wide, and distant 2.8 mm from the external edges of the sVNS electrodes. For electrode coordinates see Supplementary Table 1 and Supplementary Figure 1.

Stimulation was performed sequentially on all electrode pairs, i.e. driving current through electrode pads positioned at the same angular coordinates in both electrode rings. Stimulation was applied sequentially to all 14 pairs, using current biphasic pulse trains and stimulation/resting times with equal duration for each pair, e.g. 5s on, 5s off. For each of the target organs/functions to be identified by sVNS, a set of starting parameters was defined comprising of current pulse amplitude, width and repetition frequency. These parameters were applied for the first round of trial-and-error sVNS and modified until target response was observed on less than 50% of the electrode pairs, i.e. 7 out of 14.

Improvement of selectivity was mostly performed by adjusting pulse amplitude but in a limited amount of cases pu lse width was subject to adjustment to deliver a larger amount of charge to the nerve without compromising the electronics with large current amplitudes.

The main target function was cardiac and was monitored by recording and displaying in real time the variations in HR during the trials. Additional target functions were presence/absence of laryngeal contraction, monitored by recording laryngeal EMG, and pulmonary function, monitored by recording breathing rate during periods of spontaneous breathing. These three functions were overall assessed similarly to a previous study which performed trial-end-error sVNS on the same organs

Starting parameters for each function were:

- Cardiac (efferent, pre-vagotomy) – 1mA, 1ms, 10Hz, 15s on/off
- Cardiac (afferent, post-vagotomy) – 5mA, 2ms, 10Hz, 15s on/off
- Laryngeal – 0.2mA, 50µs, 20Hz, 5s on/off
- Pulmonary – 0.8mA, 50µs, 20Hz, 15s on/off

*2.7 Post-mortem dissection, MicroCT imaging and segmentation of vagus nerve fascicles*

#### 2.7.1 Post-mortem dissection

Following euthanasia, the sVNS cuffs were sutured to the respective nerves to ensure maintenance of position during dissection. The left and right vagus nerves were then dissected from above the nodose ganglion, across the vagotomy, and down to beyond the pulmonary branches (Fig. 2). All branches exiting the vagus within this length of the nerve were traced to their originating end-organ to verify branch type which included multiple cardiac branches (including the superior and inferior branches, and smaller branches identified near the branching point of the recurrent laryngeal branch), the recurrent laryngeal branch and a few pulmonary branches. Each sample was approximately 28 cm in length. Sutures were placed around the vagus nerve proximal to branching regions and to mark the cuff positions. The nerves were then fixed in 4% paraformaldehyde at 4°C for at least 24 hours.

**Figure 2.**
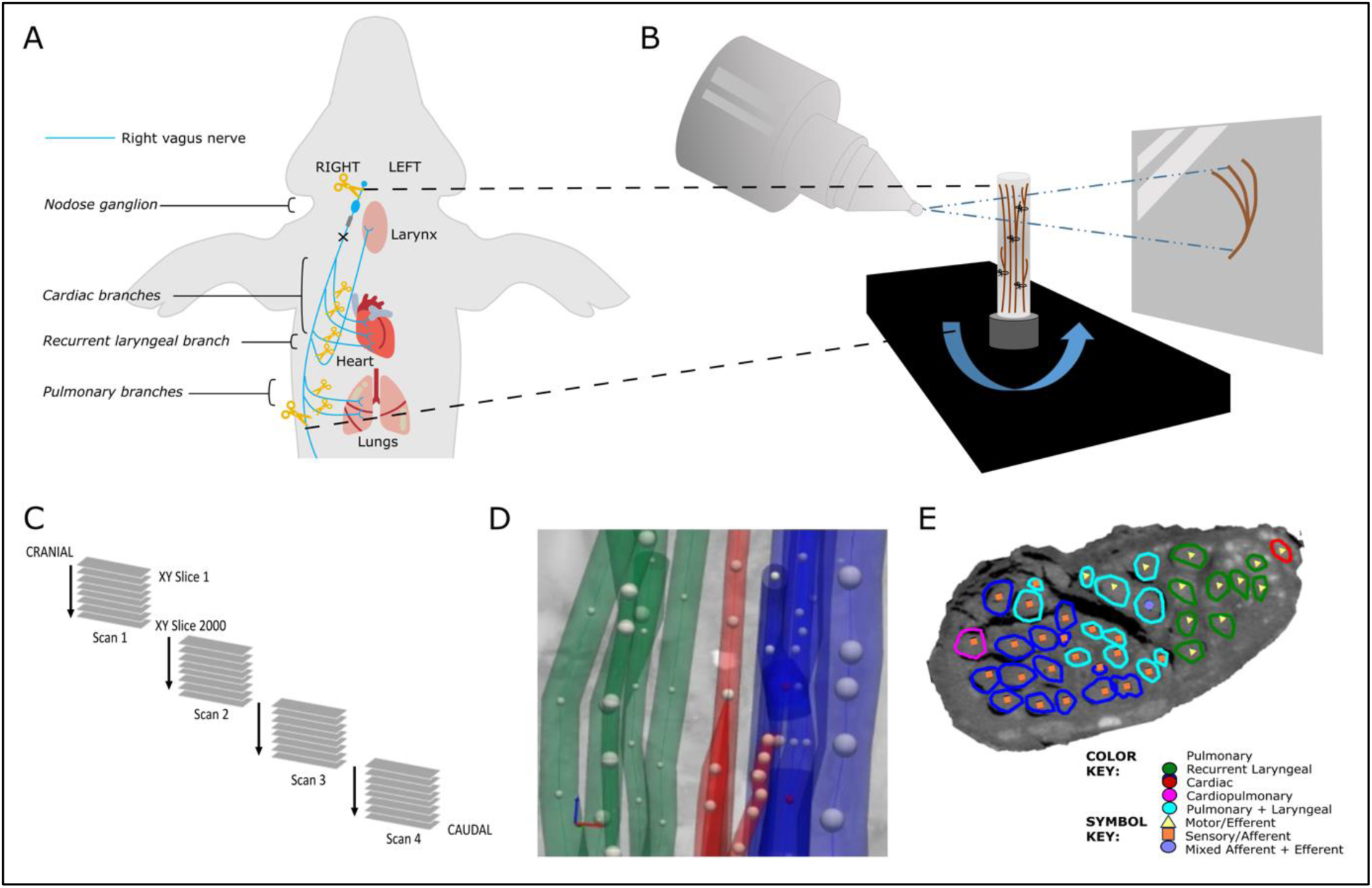
MicroCT *ex vivo* imaging. A. The right vagus of the pig is dissected post-mortem. The length of the nerve from above the nodose ganglion to below the pulmonary branches is dissected along with approximately 1cm of each branch. B. The nerve is contrast-stained, sectioned and labelled, and microCT scanned. C. Multiple, overlapping scans are required to image the full length of the bundle of nerve sections. D. The fascicles are traced from their organ origin to the cervical level by placing seed points along their path. E. The result is a labelled and mapped cross section at the mid-cervical level.

**Figure 3.**
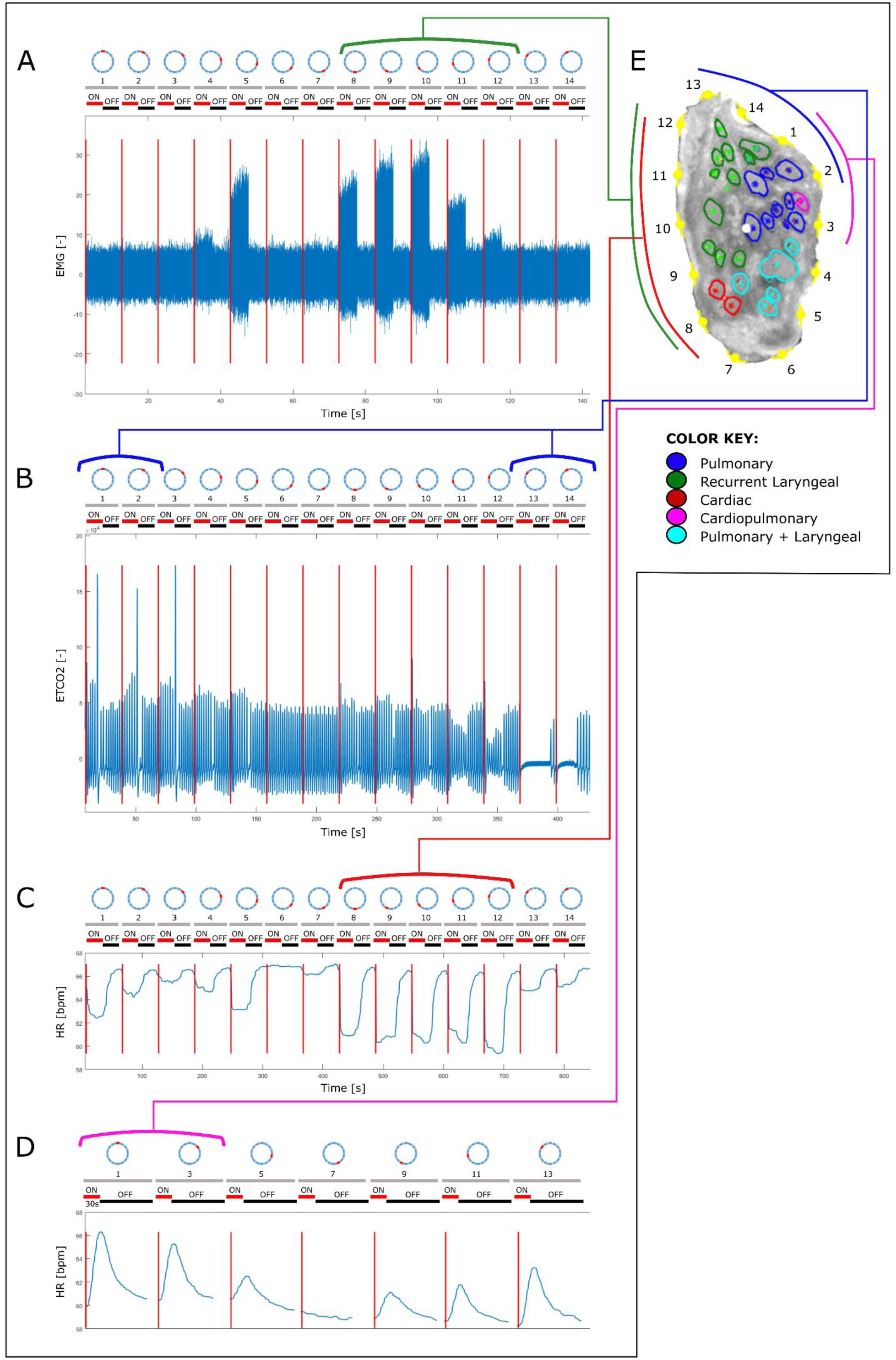
An example of the correlation between the structural and functional imaging of the porcine vagus nerve of laryngeal, pulmonary, cardiac efferent and cardiac afferent function. The structurally imaged, and microCT traced cross section (E) of the vagus nerve with electrode positions (1-14) is compared to the electrophysiological recordings of A. laryngeal EMG, B. pulmonary EtCO2, C. heart rate pre-vagotomy and D. heart rate post-vagotomy, during selective stimulation of the vagus nerve. The grey, numbered bars above each of the four traces represent the electrode positions on the sVNS cuff and the red and black bars represent the stimulation condition of on and off, respectively.

#### 2.7.2 Pre-processing, staining, microCT scanning and reconstruction

After fixation, nerve samples were measured and sutures superglued to the vagal trunk in 4.5 cm intervals, and nerves cut into 4.5 cm lengths at the level of suture placement leaving half of the suture on the end of each section as a marker for subsequent co-registration. Two to three sections were placed into a tube of 50 ml Lugol’s solution (total iodine 1%; 0.74% KI, 0.37% I) (Sigma Aldrich L6141) for five days prior to scanning to achieve maximum contrast between fascicles and the rest of the nerve tissue. On the day of the microCT scan, the nerve segments were removed from the tube, blotted dry, ordered and wrapped in cling film (10 cm x 5 cm) (Tesco, United Kingdom) in order to retain moisture during the scan. The sealed nerve samples were rolled around a cylinder of sponge (0.5 cm D x 5 cm) and wrapped in another two layers of cling film to form a tightly wound cylinder with a diameter of ∼1.5 cm to fit within the field of view at the required resolution. The wrapped cylinder was placed inside a 3D-printed mount filled with sponge around the edges, ensuring a tight fit.

A microCT scanner (Nikon XT H 225, Nikon Metrology, Tring, UK) was homed and then conditioned at 200 kVp for 10 minutes before scanning and the target changed to molybdenum. The scanning parameters were the following: 35 kVp energy, 120 µA current, 7 W power, an exposure of 0.25 fps, optimized projections, and a resolution with isotropic voxel size of 7 µm. Scans were reconstructed in CT Pro 3D (Nikon’s software for reconstructing CT data generated by Nikon Metrology, Tring, UK). Centre of rotation was calculated manually with dual slice selection. Beam hardening correction was performed with a preset of 2 and coefficient of 0.0. The reconstructions were saved as TIFF 16-bit image stack files allowing for subsequent image analysis and segmentation in various software.

#### 2.7.3 Image and nerve analysis, segmentation and tracing

Reconstructed microCT scan images were analyzed in ImageJ in the XY plane to view the cross-section of the nerve and AVI files created which allowed for validation of the scanning protocol, visual analysis of the quality of the image and the distinguishability of the soft tissue, and enabled stack slice evaluation, identification of suture positions and branching locations of the vagus nerve and served as a reference during segmentation.

Image stacks (XY plane along the Z-axis) were loaded into Vesselucida 360 (Version 2021.1.3, MBF Bioscience LLC, Williston, VT USA) and image histograms adjusted to optimize visualization of the fascicles when required. Starting from fascicle identification within branches of the vagus nerve, the fascicles were traced through the XY image stack of each scan up and through each cross section to the cervical region at the mid-level of cuff placement using the Vessel mode from the Trace toolkit. Seed points were placed in the center of the fascicle of interest along the length of the nerve at regular cross sections whilst adjusting the diameter to match the fascicle size which ultimately created a 3D segmentation of fascicles labelled according to their organ origin. Bi- and trifurcations were created when fascicles split into two or more fascicles and markers placed when fascicles merged. When fascicles labelled as containing fibers of a certain type merged with another type, the subsequent fascicle was labelled as mixed, containing fibers from both. The same process was repeated but starting at the nodose ganglion and proceeding down the nerve to mid-cervical level labelling the fascicles that bypass the nodose ganglion as efferent/motor and those that originate from the nodose ganglion as afferent/sensory. Fascicles were segmented in 2D from the rest of the nerve using the Contour mode from the Trace toolkit by forming a closed loop around the boundary of the fascicle of interest at the start or end of each scan (depending on direction of tracing) and in the mid-cuff level for visualization of the labelled fascicles in a cross section. The suture landmarks placed prior to cutting the nerve into segments were used to match up the neighboring cross sections; to continue tracing across cut regions of the nerve, the superglued suture markers and distinct physiological regions or landmarks were used to align the proximal and distal ends of the cut nerves and tracing continued.

Subsequent to the tracing of fascicles from both the proximal and distal ends, the nerves were analyzed for the number and type of fascicles at cervical level. See Supplementary Sheets 3 and 4.

### 2.8 Data co-registration and statistical analysis

See Supplementary Sheets 4 and 5 for all microCT and electrophysiological data per nerve. MicroCT data was co-registered to the circle by computing the distances from each fascicle to each of the 14 electrodes and projecting them onto the circular cross section such that radial distances to the nearest electrode were maintained. Afterwards, the area occupied by each fascicle was assigned 1, whilst remaining area was assigned 0. The resulting maps were also rotated such that the cardiac efferent fascicle was located at the top at 0 degrees for each pig (n=20 in N=5 animals, n=4 per animal).

The total “atlas” maps were computed by averaging the map for each group across all animals and normalizing it to 1. This resulted in four maps of fascicular presence across all animals, where 0 indicated that no fascicles of a particular type were present, and 1 meant that all animals had a fascicle of a particular type in a given location (N=5 animals for each map). The physiological responses recorded during selective stimulation trials were processed using back-projection mapping. First, the circular nerve cross section was subdivided onto 14 sectors, corresponding to each of the stimulating pairs. The physiological variations: HR change in % with respect to the baseline for cardiac efferent and afferent, EtCO2 change in % with respect to the baseline for respiratory, and RMS EMG change in % with respect to the baseline for laryngeal activity, were projected on every pixel within the sector of the circle for each recording in each pig (n=40 recordings in N=10 animals). The four maps (recurrent laryngeal, pulmonary, cardiac afferent, cardiac efferent) were then rotated such that the cardiac efferent area was at the top of the cross-section at 0 degrees for each pig.

Similar to microCT, the “atlas” maps were computed by averaging the map for each group across all animals and normalizing it to 1, resulting in 4 total summary maps across animals (N=10 animals for each map).

The average fascicular areas in % to the total nerve cross section for each fascicle group and technique were computed, as well as the area of overlap in % to the corresponding fascicular areas.

Centers of mass (CoMs) were computed for each fascicular group’s map for each nerve and each technique (n=40 in N=10 pigs for electrophysiology, n=20 in N=5 pigs for microCT), and the angular locations were used for ANOVA with multiple comparison tests to compute statistical significance of group localizations. This resulted in the assessment of whether the groups of fascicles/activation areas were significantly different from each other (Pulmonary vs Laryngeal, Laryngeal vs Cardiac Efferent, Cardiac Efferent vs Cardiac Afferent, Cardiac Afferent vs Laryngeal, Pulmonary vs Cardiac Efferent, Pulmonary vs Cardiac Efferent) for each technique (microCT vs selective stimulation), and if the identified locations are similar between the techniques.

## Results

### 3.1 Modulation of heart rate by selective VNS

There were 2.0±1.7 most effective electrode pairs out of 14 that elicited HR decrease at the optimal current level, which induced a -7.8±3.4% HR change, pre-vagotomy. For breathing rate (BR), the most effective pairs were 2.0±0.8 out of 14, with an induced BR change of - 73±21%. After distal vagotomy of RV, HR variations were identified by sVNS in all animals, either a decrease of -5.0±3.1% (n=4), disappearing after LV distal vagotomy or an increase of 10.2±5.3% (n=6), sustained after LV vagotomy.

The parameters of the current required to achieve spatially selective response were 1±0.2 mA amplitude, 1 ms pulse duration for activating cardiac efferent, while it was 5±3 mA, 2 ms for activating cardiac afferent fibers. For pulmonary and laryngeal selective response, the parameters were 0.8±0.2 mA, 50µs and 0.2±0.05 mA, 50µs, respectively.

A supramaximal laryngeal response was observed in every single pair for every pig while stimulating with parameters required for selective cardiac effects (all amplitudes above 1mA).

### 3.2 Composition of the vagus nerve at mid-cervical level determined by microCT tracing

The vagus nerves at the mid-cervical, mid-cuff level had a diameter of 6.63±0.6mm and an area of 2.68±0.6mm^2^. There were 1.2±0.5 cardiac, 10.2±1.8 recurrent laryngeal, 10.4±1.9 pulmonary, 1.4±0.6 cardiopulmonary, and 6±2 laryngopulmonary fiber-containing fascicles (total of 29.2±2.2 fascicles per nerve). Of these, 13.6±3.7 (47%) were identified as afferent/sensory fascicles (defined in this study as those fascicles originating from the nodose ganglion), 10.6±1.1 (36%) fascicles were identified as primarily efferent, or as those fascicles that had bypassed the nodose ganglion (henceforth referred to as efferent) and 5±3.3 (17%) fascicles were identified as containing mixed fibers. The fascicles identified as cardiac were those originating from the superior and inferior cardiac branches; these did not mix with any other fascicles within the vagus nerve. From all the nerves investigated, 100% of the purely cardiac fascicles contained only efferent fibers. Both the pulmonary and cardiopulmonary fiber-containing fascicles were identified as mostly afferent with 88.5% afferent and 9.6% mixed fibers, and 71.4% afferent and 28.6% mixed fibers, respectively. Thus, afferent projections are mixed with inputs from the heart and the lungs. Laryngeal fascicles consisted of 64.7% efferent, 17.7% afferent and 17.7% mixed fibers. Those fascicles that contained fibers originating from both the lungs and the larynx were 56.7% afferent, 13.3% efferent and 30.0% mixed (Fig. 4 and 5, Table 1, Supplementary Tables 2-5).

**Figure 4.**
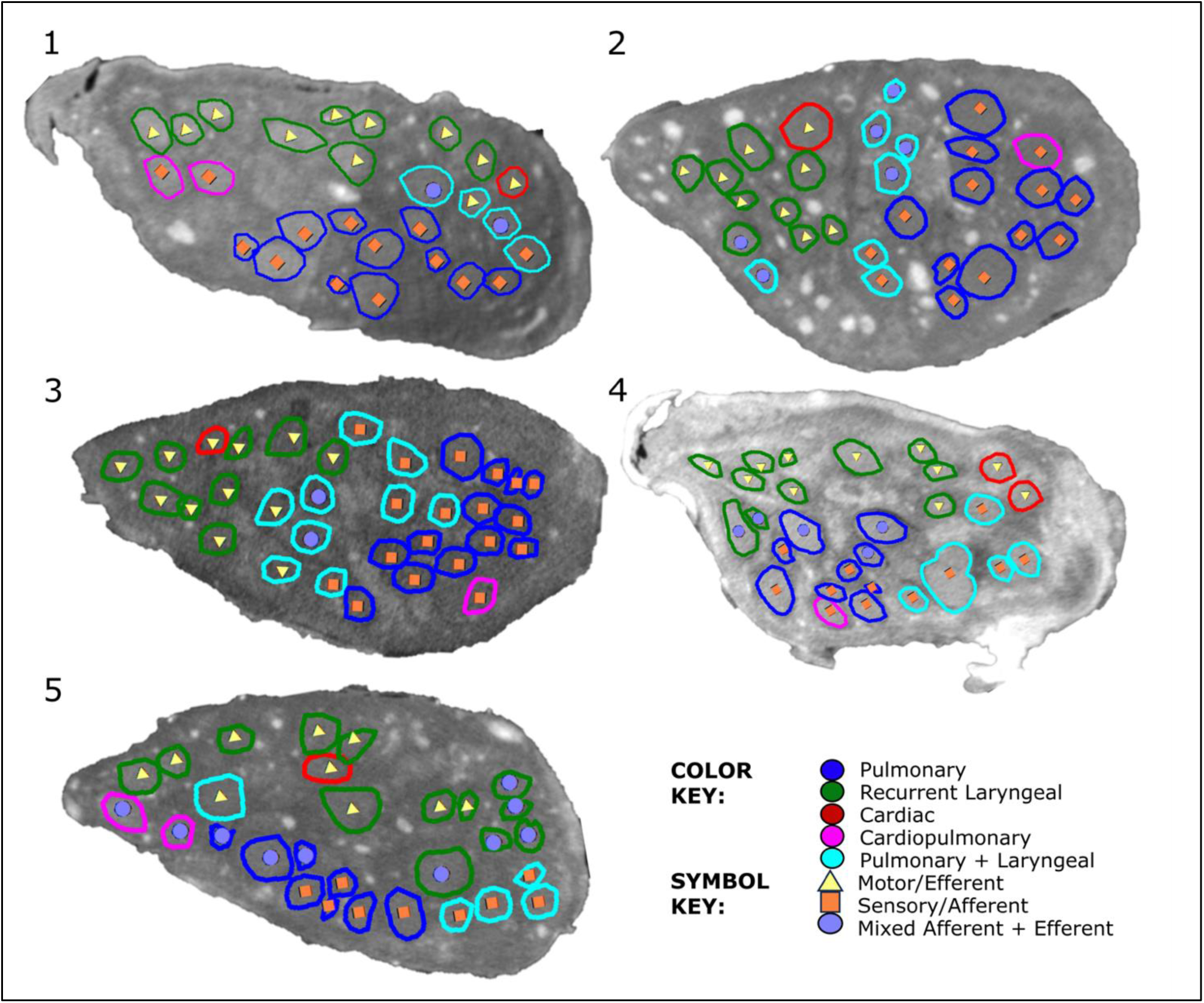
MicroCT traced and labelled nerve cross sections at the mid-cervical level for five right vagus nerves. The fascicles were traced from the three target end-organs up to the mid-cervical level to decipher the organ-specific organisation as well as down from the nodose ganglion to mid-cervical level to determine the distribution of afferent and efferent fibers.

**Figure 5.**
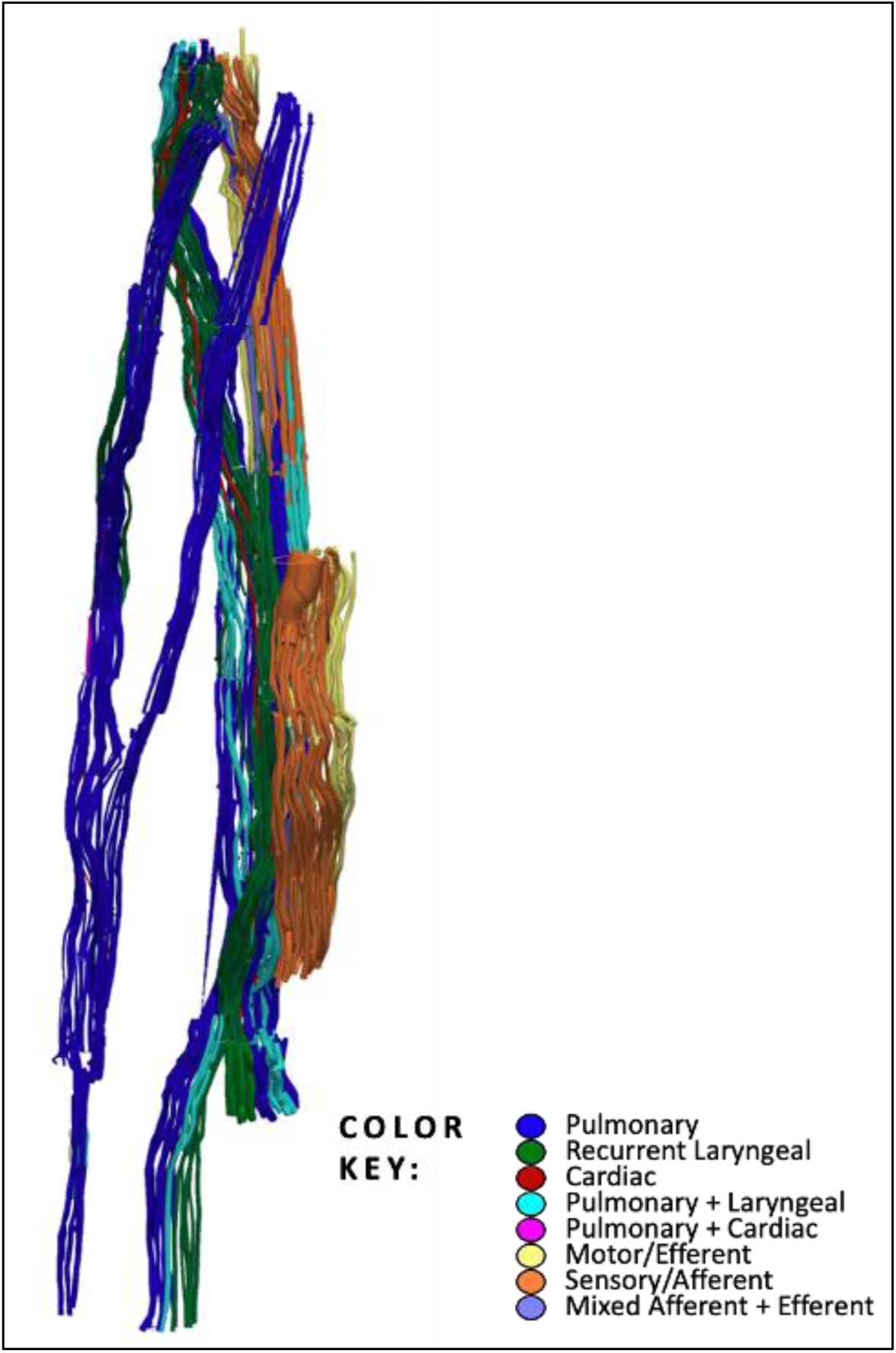
An example of 3D segmentation of the fascicles throughout the length of the vagus nerve. In order to optimise the microCT scanning of the whole length of the nerve from above the nodose ganglion to below the pulmonary branches, the nerve is cut into segments. These are bundled together within the scanner and so not all segments can be clearly visualised in this 2D example. See color key for fascicle type identification. The organ-specific (pulmonary, recurrent laryngeal and cardiac) fascicles were traced from the branches exiting the vagus up to the cervical level. The afferent and efferent-containing fascicles were traced from the nodose ganglion, and those bypassing the ganglion, down to the cervical level.

**Table 1.** Proportions and counts of organ- and fiber type-specific fascicles of the mid-cervical vagus nerve from microCT tracing data.

|  |  | *Nerve* |  |  |  |  | *Mean* | *Std Dev* | % *Total* | % *Type* |
| --- | --- | --- | --- | --- | --- | --- | --- | --- | --- | --- |
|  |  | *1* | *2* | *3* | *4* | *5* |  |  |  |  |
| *No. of fascicles* | *C* | 1 | 1 | 1 | 2 | 1 | 1.20 | 0.45 | 4.11 | 100.00 |
|  | *C - Aff* | 0 | 0 | 0 | 0 | 0 | 0.00 | 0.00 | 0.00 | 0.00 |
|  | *C - Eff* | 1 | 1 | 1 | 2 | 1 | 1.20 | 0.45 | 4.11 | 100.00 |
|  | *C - Mixed* | 0 | 0 | 0 | 0 | 0 | 0.00 | 0.00 | 0.00 | 0.00 |
|  | *L* | 9 | 9 | 9 | 11 | 13 | 10.20 | 1.79 | 34.93 | 100.00 |
|  | *L - Aff* | 0 | 0 | 9 | 0 | 0 | 1.80 | 4.02 | 6.16 | 17.65 |
|  | *L - Eff* | 9 | 8 | 0 | 9 | 7 | 6.60 | 3.78 | 22.60 | 64.71 |
|  | *L - Mixed* | 0 | 1 | 0 | 2 | 6 | 1.80 | 2.49 | 6.16 | 17.65 |
|  | *P* | 11 | 11 | 13 | 9 | 8 | 10.40 | 1.95 | 35.62 | 100.00 |
|  | *P - Aff* | 11 | 11 | 13 | 6 | 5 | 9.20 | 3.49 | 31.51 | 88.46 |
|  | *P - Eff* | 0 | 0 | 0 | 0 | 0 | 0.00 | 0.00 | 0.00 | 0.00 |
|  | *P - Mixed* | 0 | 0 | 0 | 3 | 2 | 1.00 | 1.41 | 3.42 | 9.62 |
|  | *CP* | 2 | 1 | 1 | 1 | 2 | 1.40 | 0.55 | 4.79 | 100.00 |
|  | *CP - Aff* | 2 | 1 | 1 | 1 | 0 | 1.00 | 0.71 | 3.42 | 71.43 |
|  | *CP - Eff* | 0 | 0 | 0 | 0 | 0 | 0.00 | 0.00 | 0.00 | 0.00 |
|  | *CP - Mixed* | 0 | 0 | 0 | 0 | 2 | 0.40 | 0.89 | 1.37 | 28.57 |
|  | *P+L* | 4 | 7 | 9 | 5 | 5 | 6.00 | 2.00 | 20.55 | 100.00 |
|  | *P+L - Aff* | 1 | 2 | 5 | 5 | 4 | 3.40 | 1.82 | 11.64 | 56.67 |
|  | *P+L - Eff* | 1 | 0 | 2 | 0 | 1 | 0.80 | 0.84 | 2.74 | 13.33 |
|  | *P+L - Mixed* | 2 | 5 | 2 | 0 | 0 | 1.80 | 2.05 | 6.16 | 30.00 |

### 3.3 Organization of afferent and efferent cardiac fibers in the porcine mid-cervical vagus nerve

Fascicles referred to as cardiopulmonary, identified in microCT tracing as containing fibers from both the heart and lungs, are also referred to here as cardiac afferent as they contain fibers originating from the nodose ganglion (afferent) and those going to the heart (cardiac). This corresponds with those areas of the nerves referred to as cardiac afferent from the selective stimulation data.

Co-registration of CoMs from the microCT data (n=5) of the four organ-specific regions, including laryngeal, pulmonary, cardiac efferent and cardiac afferent, showed significantly different locations within the cross section of the nerve (P<0.05) (Supplementary Tables 6-9). Analysis of the CoMs from the electrophysiology data (n=10) showed significant separation (P<0.05) between all regions of the vagus nerve cross section except for cardiac efferent and laryngeal activity (P>0.05) (Fig. 6). Between the *in vivo* and *ex vivo* techniques, the location of the cardiac efferent, cardiac afferent and pulmonary regions within the nerve was consistent i.e., there was no significant difference in the location of the CoMs (P>0.5). However, the laryngeal regions were located in significantly different areas of the nerve as determined by microCT and selective stimulation (P<0.05). This discrepancy was most probably caused by large variability and spread of laryngeal fascicles across the nerve, distortions caused by the preservation preparations, and the dominant presence of connective tissue around that area of the array.

**Figure 6.**
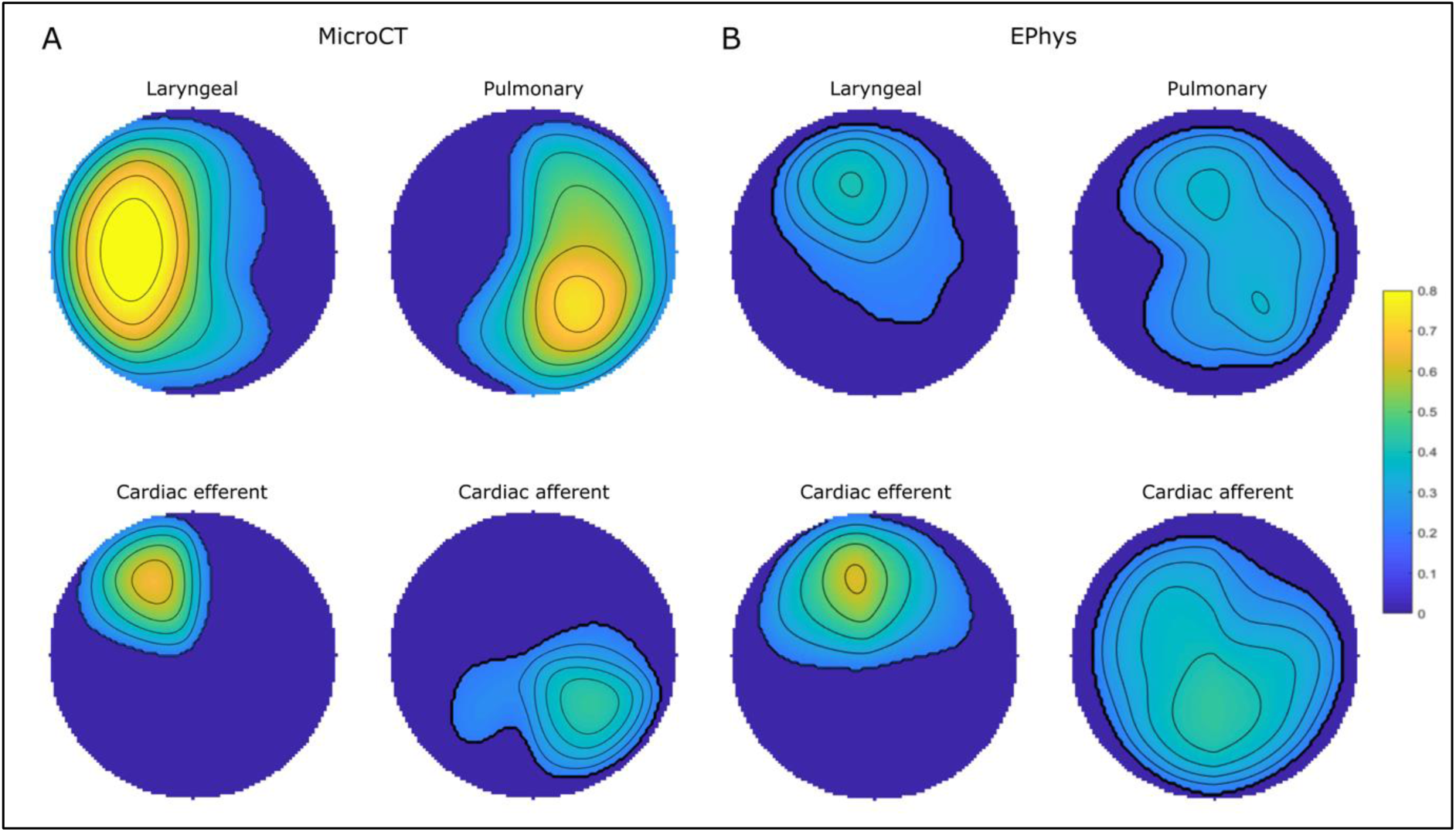
The mean regions comprising laryngeal, pulmonary, cardiac efferent and cardiac afferent fascicles or function from A. microCT (n=5) and B. electrophysiological (EPhys) data following selective stimulation (n=10), respectively, conformed to a circle for cross-validation. The colour scale, from 0 to 1, depicts the proportion of fascicles or electrophysiogical response upon selective activation that were located within the area in the cross section across all the nerves (0 indicates no fascicles were present at the location in any nerve, 1 indicates that all nerves had a specific fascicle in that location).

Notably, significantly different angular locations of the CoMs were identified for the cardiac afferent and cardiac efferent regions by microCT tracing (179±55° between CoMs around the center point of the nerve circular cross-section, p<0.05) and the post- vs pre-vagotomy cardiac neuromodulation sites identified with selective stimulation (200±137°, p<0.05).

From the fraction of the total area of nerve occupied by the respective regions identified by the two imaging techniques, there was a 0% overlap between cardiac afferent and efferent regions, a 93% overlap of cardiac afferent with pulmonary, 48% overlap of pulmonary with cardiac afferent and only a 30% and 10% overlap of cardiac efferent with pulmonary and pulmonary with cardiac efferent, respectively, identified with microCT tracing (Supplementary Table 6 and 7). With the electrophysiology data, the fractions of the areas of the nerve that overlapped between cardiac afferent and efferent, cardiac afferent and pulmonary, and cardiac efferent and pulmonary were 43/82%, 78/96%, 79/52% vice versa, respectively.

## Discussion

Overall, there was a significant spatial separation of the cardiac afferent and the cardiac efferent regions within the cross section of the mid-cervical vagus nerve at the level of VNS cuff placement. This was shown with both *in vivo* and *ex vivo* methods. Fascicles identified with *ex vivo* microCT tracing as originating from the superior and inferior cardiac branches of the vagus nerve and also identified to be fascicles that bypassed the nodose ganglion were on average on the opposite side of the nerve of fascicles identified as containing afferent (stemming from the nodose ganglion) fibers of both cardiac and pulmonary origin. In addition, cardiac recordings from the vagus nerve *in vivo* pre- (efferent) and post- (afferent) vagotomy identified significantly separated neuromodulation sites. The location of the cardiac afferent region appeared to be predominantly located within or near the pulmonary region of the nerve. The cardiac efferent regions were located in close proximity to the recurrent laryngeal regions. This is consistent with the roughly equitable spread across the nerve of the afferent and efferent fibers.

The afferent effect, determined subsequent to right distal vagotomy and thus eliminating the possibility of right efferent activity, was different across the 10 animals. The activation of the afferent fibers of the right vagus nerve either resulted in tachycardia (n=6) or bradycardia (n=4) (Fig. 7). The possibility of vagal influence on the heart via reflexively activated efferent fibers of the left vagus during stimulation of right afferent fibers was ruled out in the control recordings with right afferent vagus stimulation before and after left vagotomy. If tachycardia was achieved during afferent stimulation, this persisted subsequent to left vagotomy suggesting a reflex activation of the sympathetic chain. If bradycardia was the afferent effect achieved upon right vagus nerve stimulation, this effect was eliminated subsequent to left vagotomy confirming the effect was due to activation of the parasympathetic efferent pathway via the left vagus nerve. There was no correlation found, during these studies, between the stimulation parameters and the reflex pathway, of the left vagus nerve or sympathetic chain achieving bradycardia and tachycardia, respectively, that was activated by reflex afferent stimulation in each animal. This needs to be investigated further. As seen in Fig. 3 D, there was a delay in response upon afferent stimulation, post right vagotomy, which could be owed to the time required for reflex action via the brainstem or higher centers of the brain, and the subsequent activation of an efferent pathway. Due to the location of stimulation in the cervical neck region, it is predicted that there would be few sympathetic fibers; however, the possibility of affecting vagal sympathetic efferent function via afferent activation cannot be dismissed ^29^.

**Figure 7.**
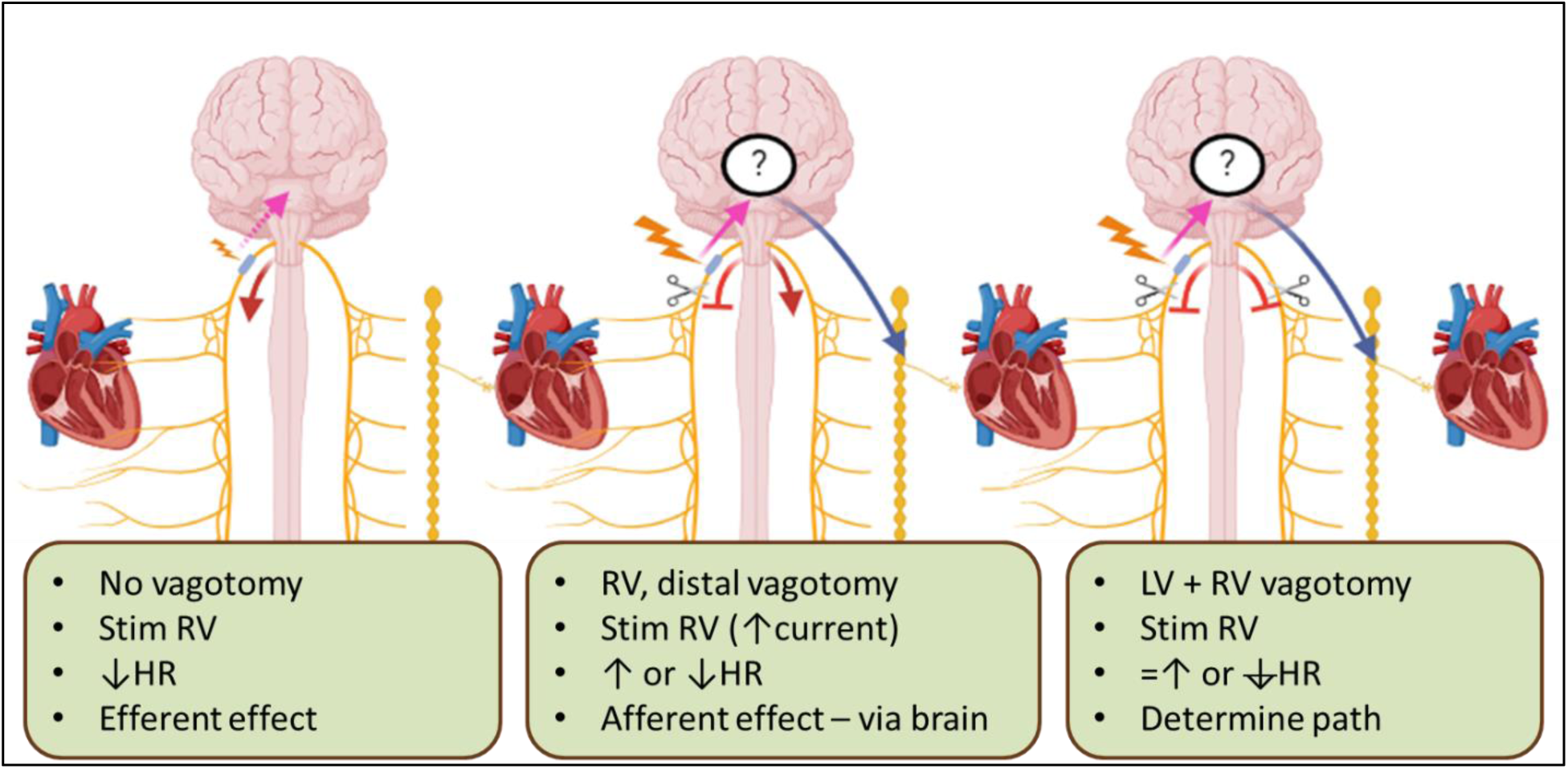
Effects of vagal stimulation on the heart at different stages in the experiments. Left: Stimulation of the right vagus nerve pre-vagotomy, achieving efferent effects (red arrow) on the heart from the right vagus nerve. Potential afferent activation (dashed pink arrow) could be possible; however, the stimulation parameters used are at a threshold lower than required to activate the afferent/sensory/smaller fibers but sufficient to predominantly activate the efferent/motor/larger fibers. See Supplementary Figure 2 for a control performed: the right vagus nerve which was activated before and after right proximal vagotomy with little change (P<0.05) in the bradycardia response. Middle: Stimulation of the right vagus nerve post right distal vagotomy, elimination of any efferent effects from the right vagus nerve (red stopper), but achieving afferent (pink arrow) effects that reflexively result in either parasympathetic efferent cardiac effects via the left vagus nerve (bradycardia, decrease in heart rate (HR)) (red arrow) or sympathetic cardiac effects via the sympathetic chain (tachycardiac, increase in HR) (blue arrow). Right: Stimulation of the right vagus nerve post right distal vagotomy and left vagotomy, resulting in the elimination of efferent effects via either vagus (2x red stoppers) which eliminated bradycardia (red stopper) or the persistence of tachycardia (blue arrow) in those animals for which bradycardia or tachycardia was the afferent effect achieved, respectively.

The development of sVNS technique could help address the complexity of performing effective neuromodulation of cardiac function, by targeting individual vagus branches or areas of the nerve responsible for cardiac control; however, a deeper understanding of the functional and anatomical organization of the various cardiac nerve fibers in the cervical vagus is required for this to be possible. Here, we showed that the cardiac efferent and afferent fibers are located separately in the cross section of the vagus nerve at mid-cervical level, which is a significant advancement in the development of sVNS approach to modulate cardiac function.

In Thompson *et al* (2023) ^20^ the fascicles of the superior and inferior cardiac branches merged proceeding up the nerve but maintained separation from the rest of the vagal fascicles at the level of vagal cuff placement at mid-cervical level. This is consistent with what was seen here in these nerves. These fascicles were identified as being efferent/motor by segmentation and tracing from microCT scans of the fascicles from those that bypassed the nodose ganglion. Majority of the fascicles containing either pulmonary fibers or pulmonary and cardiac fibers were identified as afferent/sensory with the fascicles originating in the nodose ganglion. The regions within the cross section of the nerve containing cardiac efferent and cardiopulmonary afferent fascicles were roughly on opposite sides of the nerve and this corresponded with the activity identified as efferent and afferent cardiac function, respectively, by stimulation of the nerve pre- and post-vagotomy *in vivo*.

The organization of vagal fibers related to cardiac function was also investigated in a recent study in mouse ^30^, which suggested a convergence of sensory neural pathways of cardiac and respiratory neurons involving a significant fraction of both (>30%). This is congruent with the finding of cardiac afferents overlapping with the fascicles identified *ex vivo* as containing fibers from pulmonary branches as well as those areas within the nerve identified *in vivo* as having pulmonary activity and cardiac activity post-vagotomy (and therefore, no efferent activity within the nerve). Furthermore, a recent study by Veerakumar *et al* (2022) ^31^, molecularly defined two separate cardiac circuits within mice using retrograde tracing, single-cell RNA tracing and optogenetics ^31^. These consisted of the ambiguous cardiovascular circuit which selectively innervate a subset of cardiac parasympathetic ganglion neurons and mediate the baroreceptor reflex, slowing heart rate and atrioventricular node conduction in response to increased blood pressure; and the other, the ambiguous cardiopulmonary circuit with neurons that intermingled with both the cardiovascular and also innervate most or all lung parasympathetic ganglion neurons thereby having cardiopulmonary function by innervating both organs.

These findings suggest it should be possible to selectively activate the cardiac efferent fibers of the vagus nerve whilst avoiding the cardiac afferents. However, the vagus nerve at mid-cervical level appears to mostly be grouped according to efferent and afferent on each half of the cross section of nerve. Therefore, the cardiac efferent fascicles are located near the recurrent laryngeal fascicles in all nerves in this study. Therefore, it may not be possible to avoid all laryngeal off-target effects when selectively activating the cardiac efferents as some fascicles in the vicinity may be simultaneously activated. Additionally, due to the activation threshold of cardiac fibers being 100 times greater than that for laryngeal fibers, it would be difficult to not activate recurrent laryngeal fibers simultaneously. However, this could abolish the need for titration thereby saving crucial time in treating and improving outcomes of MI and HF.

Other efforts in decoding functional signals such as cardiac- or pulmonary-related involve the use of invasive methods such as intraneural electrodes in pigs ^32^ or microelectrodes in awake humans ^33^. Machine learning has also been proposed as a potentially helpful tool for this decoding effort ^34^. Additionally, optogenetic stimulation of efferents at the level of the dorsal motor nucleus of the vagus nerve has been performed in sheep ^35^.

Recent studies pioneered the concept of neural fulcrum, based on the notion that when neuromodulation alters autonomic control, the effect is counteracted by endogenous reflexes ^10,29^. These studies also suggested the use of titration as a way to condition autonomic neural networks over time and implement stimulation protocols with reduced side effects ^10^. The ANTHEM-HF clinical study, based on the concept of neural fulcrum and titration, investigated the safety and feasibility of performing cervical VNS in patients with chronic HF and reduced ejection fraction ^36^. In a 42-months follow-up, this therapy was found to be safe and associated with beneficial effects on left ventricular ejection fraction. Titration, however, takes approximately four weeks to achieve. By selectively stimulating the cardiac efferents and avoiding majority of the afferent fibers of the vagus nerve, titration could be avoided. As shown in Hadaya *et al* (2023) ^3^, considerable cardioprotection was achieved when VNS was applied at the neural fulcrum starting two days post-MI. However, titration subsequent to MI was required to reach this optimal point. Furthermore, in Ardell *et al* (2015) ^29^, when the vagus nerve was cut centrally, the threshold for bradycardia was markedly reduced. Therefore, using a sVNS approach and activating cardiac efferent fibers whilst avoiding afferents, and thereby not having the concurrent activation of cardiopulmonary and laryngeal afferents, therapeutic levels in pure reactive mode could be achieved within days and not weeks. This could save valuable time and improve outcomes and survivability drastically. This selective neuromodulation approach holds potential for extending its therapeutic benefits to atrial fibrillation. Previous research utilizing VNS in a neutrally induced model of atrial fibrillation has highlighted its efficacy in targeting specific populations of intrinsic cardiac neurons within the atria, thereby mitigating neural network imbalances and conferring electrical stability ^25^. Considering the demonstrated effectiveness of VNS in this regard, the introduction of a selective approach could further optimize its therapeutic impact on atrial fibrillation, presenting an avenue for enhanced treatment strategies in this condition too.

While heart rate response served as the key biomarker for assessing cardiac effects following neuromodulation in this study, it is worth recognizing that the impact of neuromodulation likely extends to ventricular dynamics as well. Through comprehensive preclinical investigations, which encompassed studies utilizing atrial and ventricular indices alongside an exploration of the structure-function organization of the thoracic vagosympathetic nerve trunk, valuable insights into the multifaceted effects of neuromodulation on cardiac function were gained ^10,29,37,38^. Notably, results obtained using observed chronotropic effects are paralleled by dromotropic and inotropic effects in both atrial and ventricular tissues, highlighting the broad-reaching influence of neuromodulation on various aspects of cardiac physiology ^37,38^. Based on the observed heart rate changes in this study utilizing the novel approach of sVNS, it is anticipated that a similar correlation between control of other heart functions will persist.

This research study, despite its promising findings, possesses certain inherent limitations that need to be acknowledged. First and foremost, the high amplitudes required for successful sVNS may not be easily translatable to human applications, as the threshold for stimulation in humans might differ significantly from the observed levels. Additionally, despite bilateral vagotomy at the end of the experiment to control for left efferent input, the study did not address the potential influence of other neural circuit pathways that were left intact, such as the sympathetic chain and other spinal cord inputs. Understanding the interplay between these pathways and the effects of their preservation on the outcomes could provide a more comprehensive picture. Furthermore, the study utilized pig models, and the translatability of results to humans may be limited due to differences in fascicle number, size, and distribution between the two species. Finally, the potential contribution of co-release of neuromediators (e.g. peptides) during sVNS should be considered in addition to the primary postganglionic neurotransmitters remains to be determined. These limitations underscore the need for further investigation and refinement before the full clinical potential of sVNS can be realized.

Further elucidation of the mechanisms underlying the inconsistent effects of vagal afferent fiber activation on heart rate is warranted, particularly regarding the potential reflexive activation of sympathetic efferent pathways or left vagus nerve efferents. Given the complexities involved and the extensive scope of potential pathways and physiological factors to consider, elucidating these mechanisms falls beyond the confines of the current study. This uncertainty not only emphasizes the need for further investigation to better understand the reflex pathways in the brainstem and higher centers of the brain but it also further supports the rationale of selectively targeting efferent fibers.

Future work will encompass additional and chronic animal studies as well as future human trials, as this is essential for validating the potential of sVNS in addressing HF, MI and other applications. Our upcoming advancements in technology, including a battery-free, off-the-shelf sVNS implantable stimulator and a pressure-sensitive sVNS cuff to prevent excessive nerve pressure, will contribute significantly to enhancing the feasibility and effectiveness of sVNS for these clinical purposes.

## List of Abbreviations

VNS: vagus nerve stimulation
HF: heart failure
ANS: autonomic nervous system
MI: myocardial infarction
GI: gastrointestinal
FDA: Food and Drug Administration
sVNS: selective vagus nerve stimulation
microCT: micro-computed tomography
EIT: electrical impedance tomography
RV: right vagus nerve
LV: left vagus nerve
RL: recurrent laryngeal nerve
EMG: electromyography
AWERB: Animal Welfare and Ethical Review Body
ECG: electrocardiography
EtCO2: end-tidal carbon dioxide
HR: heart rate
EEG: electroencephalography
PEDOT:pTS: poly(3,4 ethylenedioxythiophene)/polythiophenesulfonyl chloride
RMS: root mean square
CoMs: center of mass
ANOVA: analysis of variance
BR: breathing rate

## Data availability

The data from the research in this paper will be made available on Pennsieve with the following DOI: **10.26275/0bdd-rwbe.**

